# Transmission cycles generate diversity in pathogenic viruses

**DOI:** 10.1101/339861

**Authors:** Daniel Sigal, J.N.S. Reid, L.M. Wahl

## Abstract

We investigate the fate of *de novo* mutations that occur during the in-host replication of a pathogenic virus, predicting the probability that such mutations are passed on during disease transmission to a new host. Using influenza A virus as a model organism, we develop a life-history model of the within-host dynamics of the infection, using a multitype branching process with a coupled deterministic model to capture the population of available target cells. We quantify the fate of neutral mutations and mutations affecting five life-history traits: clearance, attachment, budding, cell death, and eclipse phase timing. Despite the severity of disease transmission bottlenecks, our results suggest that in a single transmission event, several mutations that appeared *de novo* in the donor are likely to be transmitted to the recipient. Even in the absence of a selective advantage for these mutations, the sustained growth phase inherent in each disease transmission cycle generates genetic diversity that is not eliminated during the transmission bottleneck.

## INTRODUCTION

Many pathogens experience population dynamics characterized by periods of rapid expansion, while a host is colonized, interleaved with extreme bottlenecks during transmission to new hosts. The effect of these transmission cycles on pathogen evolution has been well-studied, with particular focus on long-standing predictions regarding the evolution of virulence (reviewed in Alizon *et al*. 2009), conflicting pressures of within- and between-host fitness (Gilchrist and Sasaki 2002; Coombs *et al*. 2007; Day *et al*. 2011; see Mideo *et al*. 2008 for review), or broader factors affecting the evolutionary emergence of pathogenic strains (Antia *et al*. 2003; Iwasa *et al*. 2003; Reluga *et al*. 2007; Alexander and Day 2010; see Gandon *et al*. 2012 for review).

In the experimental evolution of microbial populations, the impact of population bottlenecks has also been studied in some depth, both theoretically (Bergstrom *et al*. 1999; Wahl and Gerrish 2001; Wahl *et al*. 2002) and experimentally (Burch and Chao 1999; Elena *et al*. 2001; Raynes *et al*. 2014; Lachapelle *et al*. 2015; Vogwill *et al*. 2016). While severe population bottlenecks clearly reduce genetic diversity, the period of growth between bottlenecks can have the reverse effect: generating substantial *de novo* adaptive mutations and promoting their survival (Wahl *et al*. 2002). The survival of a novel adaptive lineage is predicted to depend not only to the timing and severity of bottlenecks, but on the details of the microbial life history and the trait affected by the mutation (Alexander and Wahl 2008; Patwa and Wahl 2008; Wahl and Zhü 2015).

The effects of transmission bottlenecks on the evolution of an RNA virus have been explicitly studied in a series of experimental papers, demonstrating that severe bottlenecks (one surviving individual) reduced fitness (Duarte *et al*. 1992) despite rapid population expansion between transmission events (Duarte *et al*. 1993). The magnitude of this effect depends on both the initial fitness of the lineage (Novella *et al*. 1995) and on bottleneck severity (Novella *et al*. 1996). In theoretical work, a model of a viral quasispecies undergoing periodic transmission events predicts that pathogens should maintain a mutation-selection balance with high virulence if the pathogen is horizontally transferred, if the bottleneck size is not too small, and if the number of generations between bottlenecks is large (Bergstrom *et al*. 1999).

Unlike the bottlenecks imposed in serial passaging, transmission bottlenecks in nature are not constrained by experimental control. Thus, key parameters such as the bottleneck size – the number of microbes initiating an infection – have proven difficult to estimate. Nonetheless experimental models (see Abel *et al*. 2015 for review), as well as recent techniques such as DNA barcoding (Varble *et al*. 2014) and sequencing of donor-recipient pairs in humans (Poon *et al*. 2016) have shed new light on this issue. In addition, we note that many human viruses – including human immunodeficiency virus, hepatitis B virus, and influenza A virus – reproduce by viral budding in the context of a potentially limited target cell population (Garoff *et al*. 1998); the survival of *de novo* mutations has not yet been predicted for this microbial life history. Thus, the effects of transmission bottlenecks on the genetic diversity of viral pathogens, that is, on the fate of de novo mutations, are as yet unknown.

In this contribution, we first develop a deterministic model of the within-host dynamics of early infection by a viral pathogen. We couple this to a detailed life-history model, using a branching process approach to follow the fate of specific *de novo* mutations that are either phenotypically neutral, or affect various life-history traits. These techniques allow us to predict which adaptive changes in virus life history are most likely to persist, and how the diversity of the viral sequence is predicted to change between donor and recipient. We can thus predict, for example, the rate at which *de novo* single nucleotide polymorphisms arise during the course of a single infection, and are transmitted to a subsequent host.

Throughout the paper, we will illustrate our results with parameters that have been chosen to model the life history and transmission dynamics of influenza A virus (IAV). IAV is an orthomyxovirus (Bouvier and Palese 2008) that imposes a significant burden on global health, causing seasonal epidemics, sporadic pandemics, morbidity and mortality (Carrat and Flahault 2007). It is estimated that infection with seasonal strains of influenza results in around 36,000 deaths per year in the United States, although exact numbers are difficult to determine (Chowell *et al*. 2008).

Mathematical modelling is a well-established tool for predicting the evolution of influenza (Larson *et al*. 1976; Bocharov and Romanyukha 1994). Because of the critical importance of immune evasion in influenza, interest has focused on the adaptation of the virus in response to immune pressure, focusing on antigenic drift (Boianelli *et al*. 2015) and antigenic shift (Feng *et al*. 2011) in the global influenza pandemic (van de Sandt *et al*. 2012). Recent models, however, have specifically addressed the within-host dynamics of influenza A virus (Beauchemin *et al*. 2005; Baccam *et al*. 2006; Beauchemin and Handel 2011; Smith and Perelson 2011; Dobrovolny *et al*. 2013; Boianelli *et al*. 2015). In concert with these contributions, recent empirical work has elucidated the life history of the influenza A virus, providing quantitative estimates of parameters such as the minimum infectious dose (Varble *et al*. 2014; Poon *et al*. 2016), the size of the target cell population, and the kinetics of viral budding (Baccam *et al*. 2006; Beauchemin and Handel 2011; Pinilla *et al*. 2012). Although we now have an increasingly clear picture of the within-host life history of this important pathogen (Beauchemin and Handel 2011; Biggerstaff *et al*. 2014), estimates of the rate at which *de novo* mutations arise and are transmitted have not yet been available. Our approach allows direct access to this question.

## LIFE HISTORY AND TRANSMISSION MODEL

### Deterministic Model

We use a system of ordinary differential equations (ODEs) to approximate the within-host dynamics during the early stages of infection by a pathogenic virus, assuming a life history that involves infection of a target cell, an eclipse phase, and finally an infectious stage. Specifically, we propose:

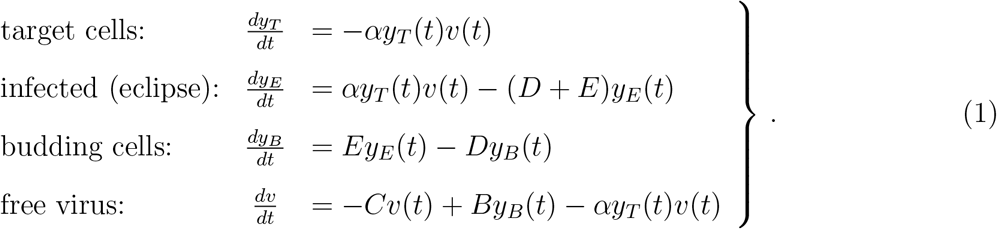

Here *y_T_* represents susceptible target cells (in the case of influenza A virus we consider epithelial cells of the upper respiratory tract), *y_E_* represents cells that are infected by the virus but not yet in the budding stage, *y_B_* represents mature infected cells (infected cells that are budding), and *v* represents the free virus, that is, virions not attached to target cells (Baccam *et al*. 2006). Parameter B gives the rate at which budding cells produce infectious free virus; *C* gives the clearance rate for free virus. Infected cells die at constant rate *D*, while *E* represents the rate at which infected cells mature, leaving the eclipse phase and becoming budding cells. The parameter *α* gives the rate of attachment per available target cell. Thus the overall attachment rate for a virion is a function of the time-varying target cell population, and can be written *A*(*t*) = *αy_T_*(*t*), with the corresponding mean attachment time, *A*(*t*)^-1^.

A limitation of ODE approaches is that all transitions are described by exponential distributions. To relax this assumption, we introduce a sequence of *k* infected stages through which infected cells pass before reaching the budding stage. This ‘chain of independent exponentials’ allows for more realistic gamma distributions of eclipse times (Wahl and Zhu 2015). Specifically, we replace system (1) with:

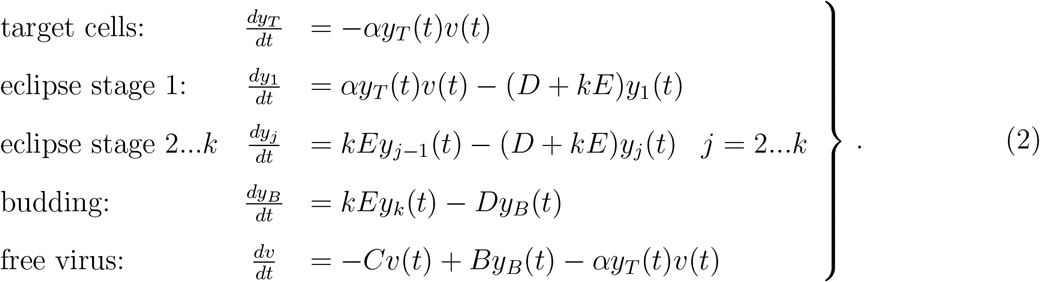

When *k* =1, this model reduces to System 1; for *k* > 1, *y*_1_ gives the population of initially infected cells, which pass through *k* eclipse stages at rate *kE* before budding. The transition rate *kE* is set such that the expected time in the eclipse phase, in total, is fixed at 1/*E* for any value of *k*. In the supplementary material, we also investigate a model in which the death term, *D*, is set to zero during the eclipse stages and only acts during the budding stage. This likewise gives a more realistic distribution for the lifetime of infected cells.

The founding virus begins as an initial population of free virus (the initial infectious dose, *v*(0) = *v*_0_) at time *t* = 0. We do not assume that all viral particles in the founding dose are genetically identical, but we do assume that they are phenotypically identical, that is, they are described by the same parameter values in the deterministic model. As described further in the stochastic model below, we assume that disease transmission occurs at time *τ* during the peak viral shedding period (when the free virus population, *v* is at or near a peak value). For the transmission event to a new susceptible individual, a new founding population is sampled from the total viral load. In particular, each free viral particle becomes part of the infectious dose transmitted to the next individual with probability *F*. The value of *F* is computed such that for the founding virus, the expected size of the transmitted sample is *v*_0_, that is, *F* = *v*_0_/*v*(*τ*). Note that only free virions – those not yet attached to a target cell – are transferred to the next individual during transmission.

Immune responses clearly play a critical role in the within-host dynamics of viral infections, as well-documented for models of influenza A (Beauchemin and Handel 2011; Smith and Perelson 2011; Dobrovolny *et al*. 2013). In the model proposed above, innate immune mechanisms are included in the clearance rate of free virus and the death rate of infected cells. Because we use this model only until the time of peak viral shedding, which occurs 54.5 hours post infection (see parameter values, below) and before the adaptive immune response is activated (Tamura and Kurata 2004), we do not include the adaptive immune response. We address this issue further in the Discussion. Likewise, we do not include replenishment of the susceptible target cell population over the initial 54.5 hours of the infection. This is consistent with complete desquamation of the epithelium (loss of all ciliated cells) within three days post-infection in murine influenza, followed by regeneration of the epithelial cells beginning five days post-infection (Ramphal *et al*. 1979).

### Stochastic Life History Model

To describe the lineage associated with a rare *de novo* mutation, a stochastic model is required. To gain tractability, we assume that the mutant lineage propagates in an environment for which the overall dynamics of the target cell population are driven by the deterministic system (2). Thus we treat the free virus, eclipse-phase cells and budding cells in the mutant lineage stochastically, but use the deterministic system to predict the susceptible target cell population at any time.

As in the deterministic model, free virions clear at a constant rate *C* or adsorb to susceptible host cells at rate *A*(*t*). Note that the attachment rate of a free virion is not constant; it depends on target cell availability, such that *A*(*t*) = *αy_T_*(*t*), where *y_T_*(*t*) is the target cell population predicted by system (2). Host cells enter the eclipse phase when a virion adsorbs, and exit the eclipse phase at rate E. After the eclipse phase, mature infected cells bud virions at rate *B*. Since budding itself does not immediately kill the host cells (Garoff *et al*. 1998), after infection the cell is subject to a constant death rate *D*, or in other words the cell remains alive for an average time 1/*D*.

This stochastic growth process can be described as a branching process, using a multitype probability generating function (pgf) to describe a single lineage of free virions (associated with dummy variable (*x*_1_), infected cells (*x*_2_), and mature cells (*x*_3_). As derived in the Appendix, the pgf for this process, *G*(*t, x*_1_, *x*_2_, *x*_3_), satisfies:

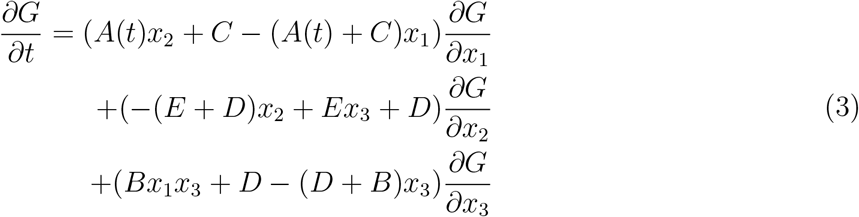

where *A*(*t*), *B, C, D* and *E* are attachment, budding, clearance, cell death and eclipse maturation rates, respectively. Equation (3) captures the time evolution of the pgf, given each of these probabilistic events. As shown in the Appendix, Equation (3) can be converted to a system of ODEs using the standard method of characteristics, and is thus amenable to numerical solution. Analogous to System (2), Equation (3) can also be extended to include a chain of *k* infected stages before the budding stage, yielding more realistic distributions of the eclipse time.

To estimate the probability that the lineage associated with a *de novo* mutation does not survive the transmission bottleneck, we will need a pgf describing a complete cycle of in-host growth followed by a transmission bottleneck. We thus numerically integrate the pgf G, described above, from time 0 to time *τ*, and then compose it with a pgf describing disease transmission. To describe disease transmission, we simply assume that each free virion in the infected host is transmitted with fixed probability *F*, as described above. As derived in the appendix, this approach allows us to estimate the probability that a *de novo* mutation that first occurs at time *t*_0_ is transmitted to the next host, 1 – *X*(*t*_0_), the rate at which such “surviving” mutations arise at each time during the infection, *v*(*t*_0_), and, ultimately, the probability that a given mutation occurs *de novo* and is transmitted to the next host, 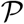.

### Beneficial Mutations

Our goal is to predict the fate of mutations that may arise *de novo* in the viral population. Although most mutations will be deleterious, we note that the virus population grows by several orders of magnitude (possibly up to seven) during a single infection, and thus deleterious mutations should be effectively purged by selection. We therefore focus in this contribution on neutral mutations (no phenotypic effect), or rare mutations that confer an adaptive advantage to the virus. For a budding virus, changes in five life history traits can confer a selective advantage: a reduction in either the cell death rate, 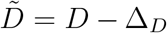, or clearance rate, 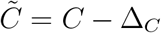; an increase in the attachment rate, 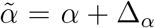, or budding rate, 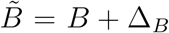; or an increase in the rate at which cells mature and begin budding, 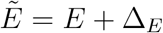.

To estimate the probability that a beneficial mutation ultimately survives, we substitute the parameters above for the analogous parameters in the pgf *G*(*t, x*_1_, *x*_2_, *x*_3_) and numerically evaluate *G*(*τ, x*_1_, 1, 1), which describes the distribution of free virions in the mutant lineage at time *τ*, as described in the Appendix. We then compose this function with the pgf describing disease transmission. The accuracy of these numerical solutions was verified using an individual-based Monte Carlo simulation, developed for a reduced model without target cell limitation, similar to the approach described by Patwa and Wahl (2009).

### Selective Advantage

Finally, in order to compare the fitness of mutations affecting different traits, we calculate the selective advantage of each mutation. Following common experimental practice, we define fitness in terms of the doubling time, that is, we assume that in the time required for the founding population to double, the mutant lineage grows by a factor of 2(1 + *s*). Given the founding growth rate *g*, we substitute the founding doubling *t* = ln(2)/*g* into 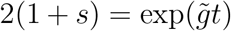 to find the selective advantage of the mutant, 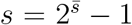, where 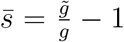. (For the relatively small *s* values presented here, this definition of the selective advantage differs from the more appropriate but less commonly used 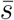 by a constant factor of ln 2.)

To estimate the average growth rates, *g* and 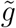, we consider a single cycle of growth, starting from a single free virus at time 0. In this case the partial derivative of *G* with respect to *x*_1_, defined as *Z* = *∂G*(*τ*, 1,1,1)/*∂x*_1_, gives the expected number of free virions at time *τ*, illustrated here for the case *k* =1 (Grimmett and Welsh 2014). The derivative was calculated numerically, and the average exponential growth rate of the free virus population is then given by *g* = ln *Z*/*τ*.

### Parameter values for influenza A virus

Parameter values were estimated where possible from the empirical and clinical literature for influenza A virus, and are displayed in Table 1. Beauchemin and Handel 2011 give a range of values for several relevant parameters, from which parameter estimates for *C, D*, and *E* were chosen. Specifically, we take the clearance time to be 3 hours, the cell death time 25 hours, and the eclipse time 6 hours (Baccam *et al*. 2006; Beauchemin and Handel 2011).

**Table 1:**
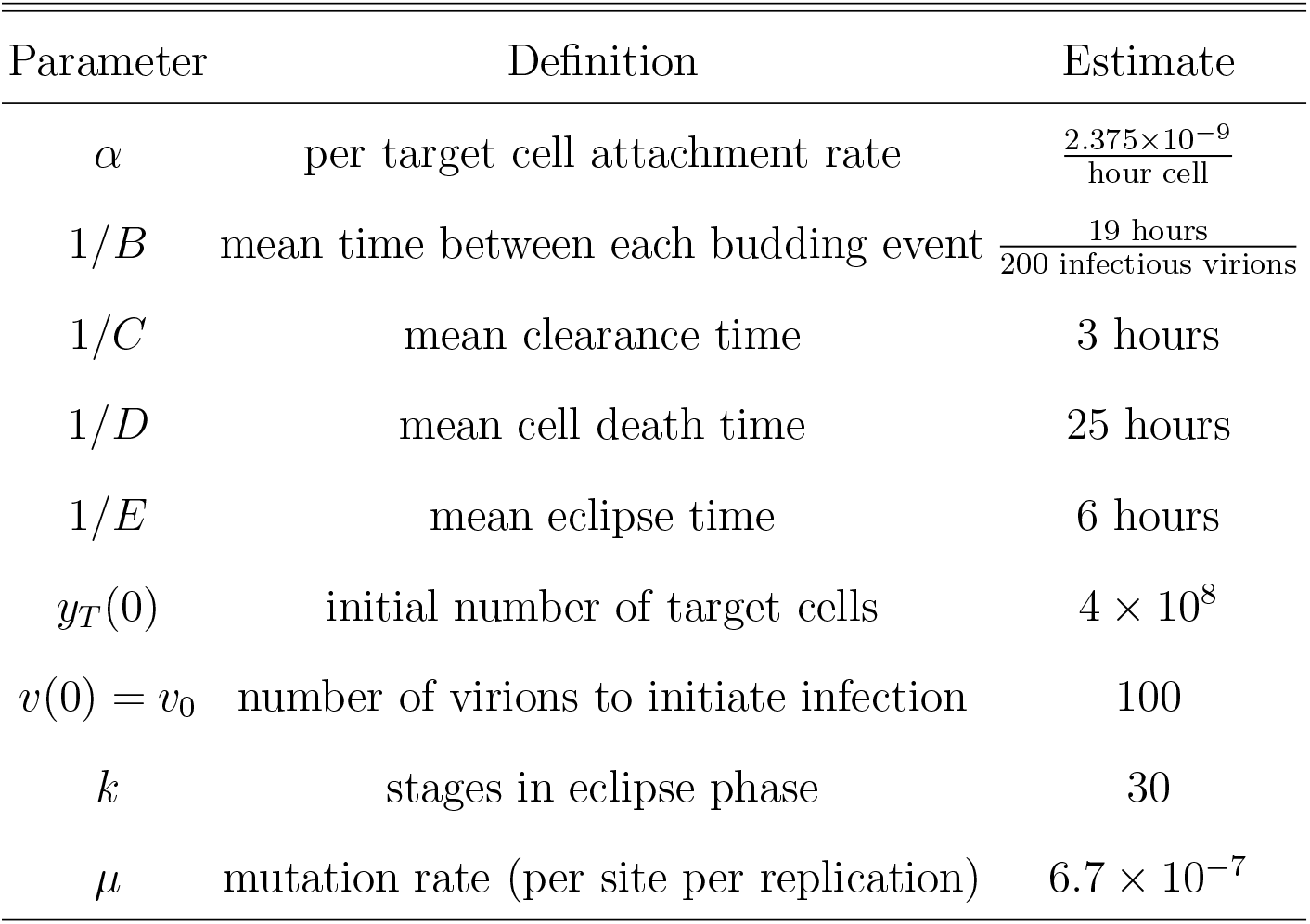
Parameter Estimates for Influenza A Virus

To estimate the time between each budding event, 1/*B*, we first consider the total number of virions produced per cell, the “burst size”. For influenza A virus, the burst size has been estimated to be between 1000-10000 virions (Stray and Air 2001). However, not all virions produced are infectious and in fact a large fraction are unable to infect a host cell; the particle to infectivity ratio for influenza A is approximately 50:1 (Martin and Helenius 1991; Roy *et al*. 2000). Taking the upper bound of the range for burst size, of the 10000 virions produced only 200 are predicted to be infectious. Recall that budding does not kill the host cell, therefore budding time depends on the eclipse and cell death times. An eclipse time of 6 hours and a cell death time of 25 hours gives a budding time of 19 hours. Therefore, the time between each infectious budding event, 1/*B*, is assumed to be assumed to be 19/200 hours per infectious virion.

The number of upper respiratory epithelial cells in a healthy adult is estimated to be 4 × 10^8^ (Baccam *et al*. 2006). Consistent with the complete desquamation of the epithelium observed in murine influenza (Ramphal *et al*. 1979), we therefore take *y_T_*(0) = 4 × 10^8^. In the supplementary material we investigate the sensitivity of our main results to this value. Similarly, we assume that an infection is founded by v0 = 100 virions, consistent with recent sequencing of donor-recipient pairs (Poon *et al*. 2016). However since values of 10-200 have been suggested in the literature (McCaw *et al*. 2011; Varble *et al*. 2014; Peck *et al*. 2015), we will also demonstrate results over a range of *v*_0_ values.

To allow for realistically distributed eclipse times, we assume a gamma-distributed eclipse phase by including a sequence of *k* infected stages before the budding stage. As described above, the mean eclipse time, 1/*E*, is set to 6 hours. The variance of the eclipse period of influenza A can then be used to estimate *k*. Pinilla *et al*. (2012) used a best-fit analysis for kinetic parameters of influenza A to predict a mean eclipse time of 6.6 hours, with an eclipse period standard deviation, *σ*, of 1.2 hours. Since the standard deviation for a gamma distribution with mean m is given by 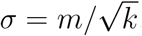, these values suggest that a realistic value of *k* is approximately 30.

We fix the attachment rate, *α*, such that that the peak of the free viral load occurs within the reported range for influenza A of 48 to 72 hours post-infection (Wright *et al*. 2001; Lau *et al*. 2010). The attachment rate *α* = 2.375 × 10^−9^ per hour per cell provided in Table 1 yields a peak time of *τ* = 54.5 hours, and implies a mean attachment time, 1/*A*(0), of just over one hour when target cells are plentiful. We assume that disease transmission is most likely at the peak viral shedding time, and thus study a transmission event that occurs at this peak time, *τ*. Note that when we examine the sensitivity of the model, for example when changing *v*_0_, we leave the attachment rate *α* fixed. We recompute the time course *v*(*t*) and assume that the transmission event occurs at the peak value of *v*(*t*). The transmission time, *τ*, then differs slightly between cases. In no case was τ outside the empirically estimated range of 48-72 hours.

The probability that each free virion survives the bottleneck and is transmitted to the next susceptible individual is defined as *F*. This probability is calculated by using the peak number of free virions, *v*(*τ*), found by numerically solving model 2. As only free virions contribute to the infectious dose, the fraction of free virions surviving the bottleneck is *F* = *v*_0_/*v*(*τ*), where again *v*_0_ is the founding population size for the next infected individual.

The mutation rate for influenza A, per nucleotide per replication, has been estimated as *μ* = 2 × 10^−6^ (Nobusawa and Sato 2006). This estimate was obtained for the IAV nonstructural gene during plaque growth, and thus does not include lethal mutations. Neglecting differences in transition and transversion rates, we divide this value by three to estimate the rate at which a specific, non-lethal nucleotide substitution occurs. We investigate the sensitivity of our results to this parameter as well.

### Data Availability

The authors affirm that all data necessary for confirming the conclusions of this article are represented fully within the article and its tables and figures.

## RESULTS

Figure 1 illustrates the deterministic dynamics of System 2, showing the time course of the inhost influenza A infection. The free virus peaks at 54.5 hours, just after the peak in the mature (budding) cell population. Note that in this simplified model, the availability of target cells limits the infection. As described earlier, this model is only accurate while the adaptive immune response remains negligible; although we illustrate the full seven days of infection, we use only the first 54.5 hours in the subsequent analysis.

**Figure 1:**
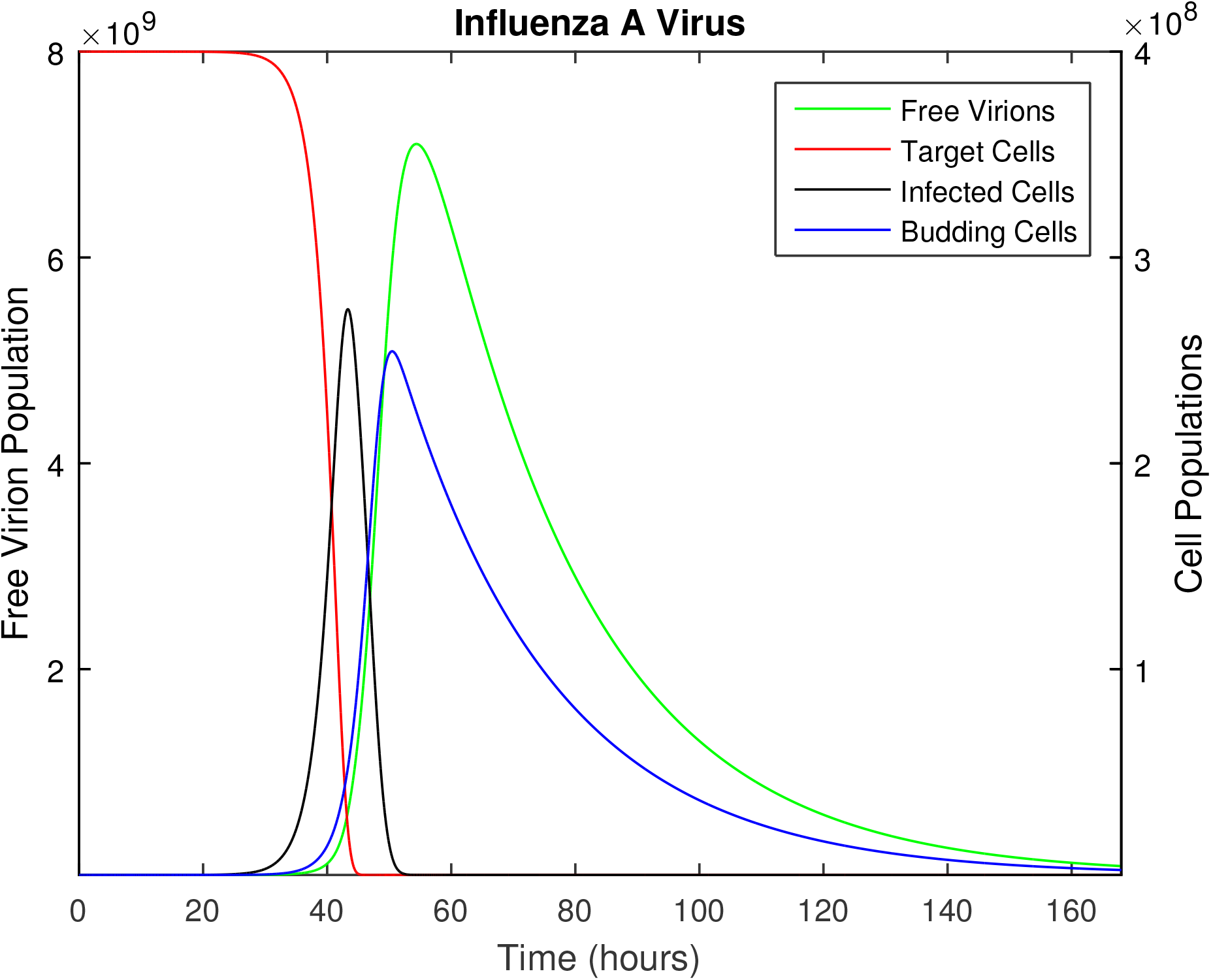
The time course of influenza A infection over the span of a week (168 hours). Parameter values are provided in Table 1, with the following initial conditions: 4 × 10^8^ epithelial cells (target cells), 100 virions (initial infection dose), all other populations initially zero.

Figure 2 shows what we will refer to as the *mutation transmission rate*, that is, the probability that at least one copy of a specific mutation arises *de novo* during an infection time course, survives genetic drift and is successfully transmitted to the subsequent host. Model predictions for beneficial mutations affecting each life history trait are shown versus the selective coefficient, s; the intercept at *s* = 0 shows the prediction for neutral mutations. Here we have assumed for comparison that the baseline mutation rate is equal for all types of mutation, however the y-axis in Figure 2 scales approximately linearly with *μ*. In the Supplementary Material, we illustrate results for a wide range of mutation rates.

**Figure 2:**
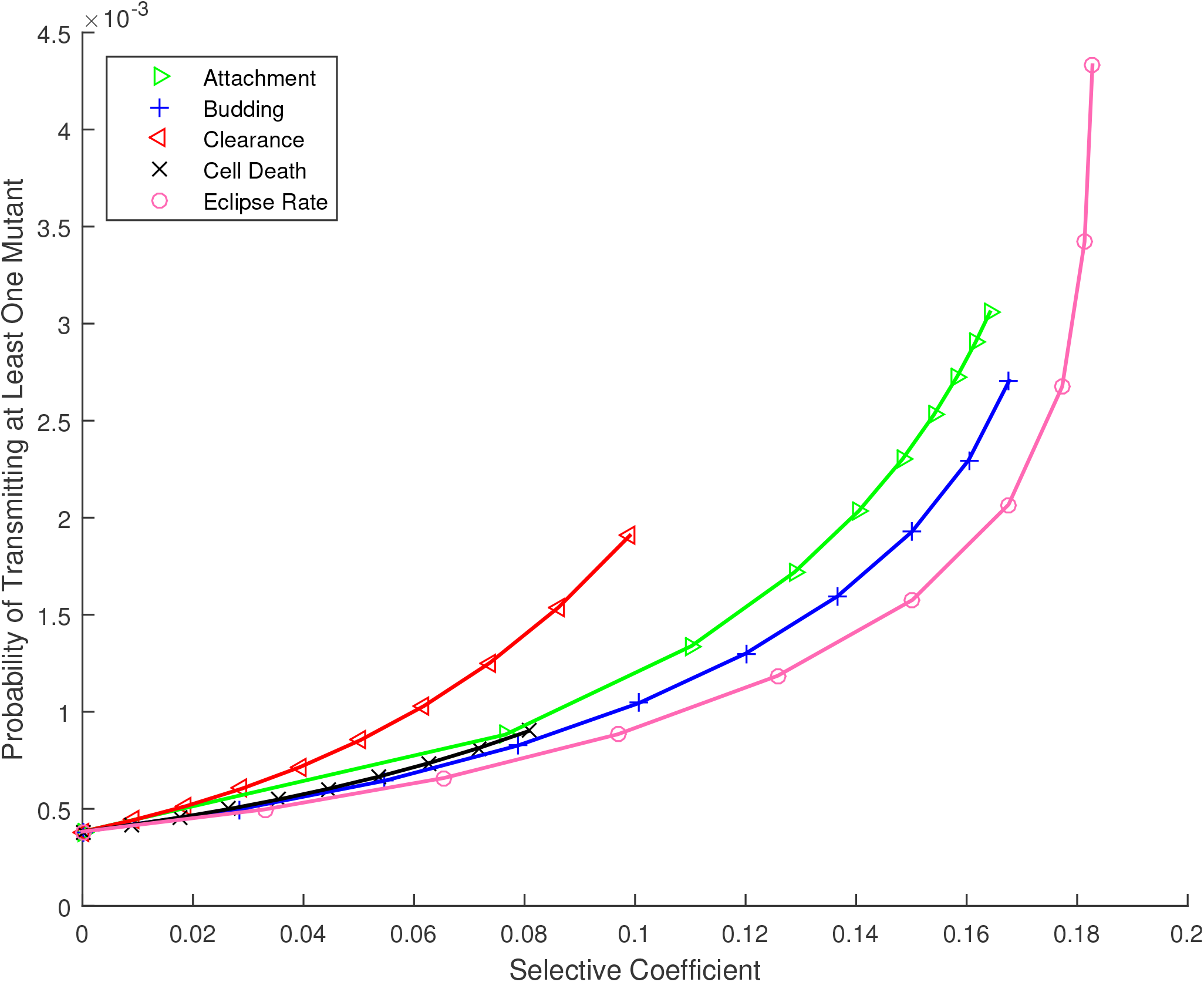
Probability that at least one *de novo* mutation arises during the infection time course and is passed to the next host, for mutations affecting the life history of influenza A virus, versus their selective coefficient. Parameters as provided in Table 1.

To interpret these results, the empirical mutation rate must be carefully considered. The rate estimate we use reflects the probability, per replication, that a specific substitution occurs at a specific nucleotide in the influenza A sequence, given that the substitution is non-lethal. Thus for example if the substitution of interest is neutral or effectively neutral, the model predicts that this substitution would occur *de novo* in the donor and be transmitted to a recipient about once in every 2000 transmission events. If the substitution of interest confers a selective advantage, the mutation transmission rate would be higher. Clearly, a large fraction of viable mutations will be deleterious and would be outcompeted before transmission; this would correspond to a lower overall mutation rate as examined in the Supplementary Material and outlined further in the Discussion.

The most striking result of Figure 2 is the predicted evolvability of influenza A during a single transmission cycle. The mutation transmission rate of one in two thousand, per substitution per site, may contribute substantial diversity since the influenza A genome is a sequence of over 13,000 nucleotides with three possible substitutions per site. We will return to the interpretation and implications of this prediction in the Discussion.

The near-overlapping lines in Figure 2 indicate that the mutation transmission rate does not vary widely across life history traits, and also illustrates the maximum selective advantage made possible by improvements to each trait. For example, clearance and cell death rates can only be reduced to zero, limiting the range of s for these traits. Although there is no upper bound on the rates of attachment or maturation to budding (eclipse rate), once these rates are effectively instantaneous, further increases do not appreciably change the growth rate, and so higher s values are also inaccessible for these traits. Similarly, increases to the budding rate cannot improve the growth rate without bound, due to target cell limitation.

Results in Figure 2 assume the default parameter set (Table 1); in particular, 100 virions are chosen at random from the free virus population and transmitted to the new host. In Figure 3, we fix the elective coefficient (*s* = 0.05) but vary the size of this transmission bottleneck. We find that the mutation transmission rate increases roughly linearly with bottleneck size.

**Figure 3:**
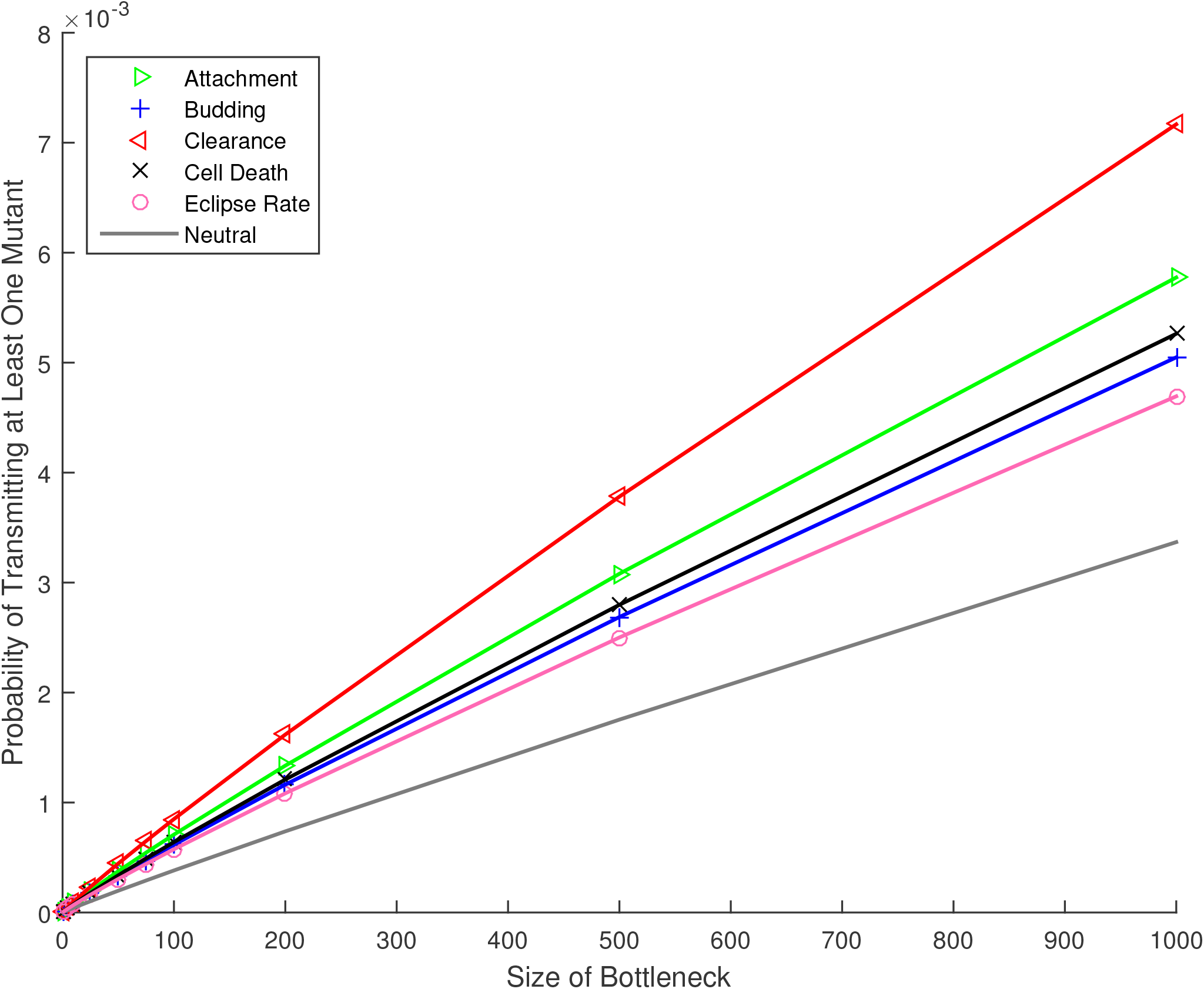
Probability that at least one *de novo* mutation arises during the infection time course and is passed to the next host, for mutations affecting the life history of influenza A virus, versus the number of virions in the transmission bottleneck. All mutations have a selective advantage of *s* = 0.05, except for the curve marked “neutral”, for which *s* = 0. Other parameters as provided in Table 1.

The results above compare mutations that have equivalent effects on the overall growth rate of the virus, assuming that the underlying mutation rate is the same for all mutations. Although the question of mutational accessibility is beyond our focus, some sense of the degree to which these mutations might be physiologically achievable can be obtained by considering the relative changes required to the trait value. To this end, Figure 4 shows the relative change in each life history parameter necessary to achieve a specific increase in growth rate (selective coefficient). To incur an advantage of s = 0.08, for example, requires less than a 10% change in the rate at which cells leave the eclipse phase and begin budding; in contrast the attachment rate would need to double (change by over 100%) to achieve the same selective advantage. Note again that clearance and cell death rates can only be reduced by at most 100%, limiting the range of their possible effects. For the other three traits, as described previously, beneficial mutations can produce selection coefficients in the approximate range 0 < *s* < 0.2, but further rate increases produce diminishing returns and fitness saturates.

**Figure 4:**
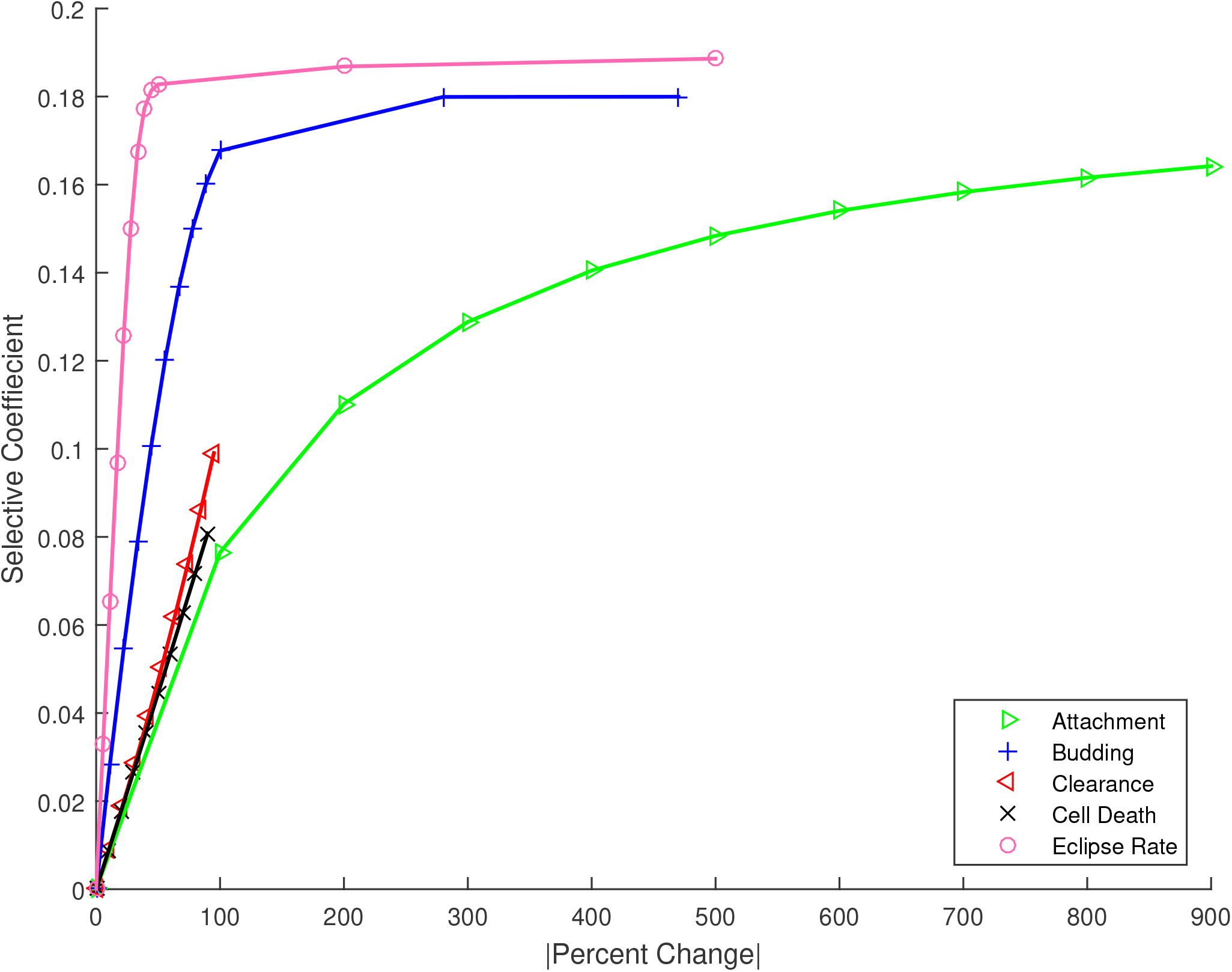
The change in selective coefficient achieved by a given absolute percent change in trait value, for mutations affecting the five life-history traits. For example, large changes in attachment rate would be required to achieve the same advantage as relatively small changes in eclipse timing.

Figure 2 gives the overall probability that a *de novo* mutation is generated and passed on. As described in the Methods, this value reflects the integrated probability of occurrence and survival for mutations that could first occur at any time during the infection time course. To better understand the dynamics of this process, in Figure 5 we show the predicted survival probability, the probability that the mutation survives and is transmitted to the next host, for mutations that arise at time *t*_0_ during the infection time course. Despite the fact that transmission to the next host occurs at 54.5 hours, the figure gives the impression that mutations that arise after about the first 10 hours of infection have little chance of survival.

**Figure 5:**
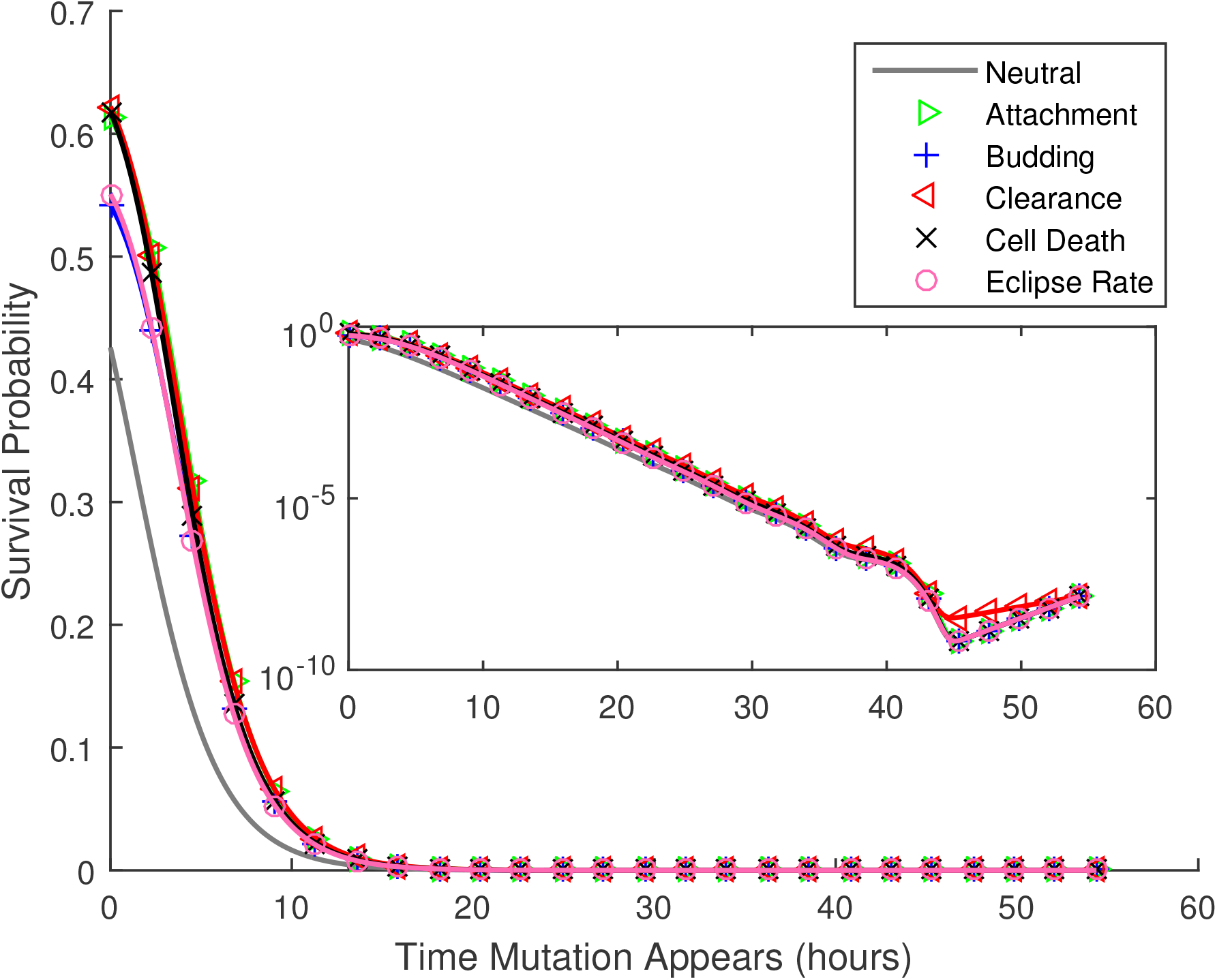
Given that a *de novo* mutation first occurs at time *t*_0_ after the start of the infection, the probability that at least one copy of it is transmitted to the next host, versus *t*_0_. All mutations have a selective advantage of *s* = 0.05, except for the curve marked “neutral”, for which *s* = 0. Parameters as provided in Table 1. The inset shows the same results with in a semilog plot.

The results in Figure 5 are mitigated, however, by the fact that many more replication events occur later during the growth phase. To investigate the rate at which surviving mutations (mutations that are transfered to the next host) first occur, we consider the product of the transmission probability for mutations that arise at each time and the number of new virions produced at that time, *By_B_*(*t*_0_).

Figure 6 shows these results. The model predicts that transmitted mutations occur throughout the infection time course, except during the first few hours of infection, when very few new virions are produced, and for a brief window approximately 10 hours before the transmission event. The latter effect presumably occurs because virions produced in this window are unlikely to be free at the time of transmission (infected cells are not transmitted). The oscillations in these curves occur because the founder virions start syncronously at *t* = 0 as free virions, and must attach and complete the eclipse phase before new virions can be produced.

**Figure 6:**
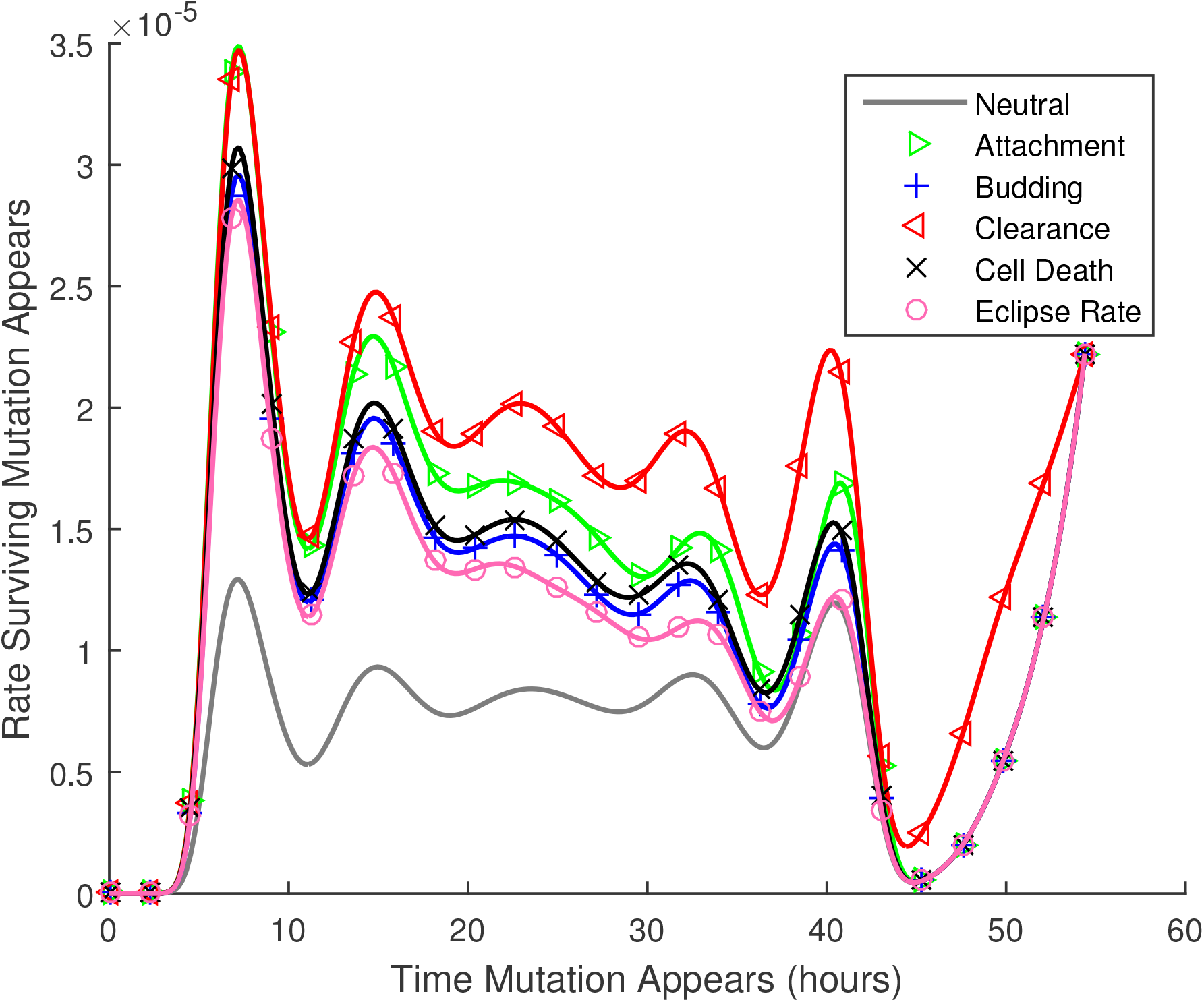
The rate at which transmitted mutations appear, versus the time at which they first appear. For this figure, the probabilities of being passed on to the next host illustrated in Figure 5 are multiplied by the number of new virions produced at each time, *μBy_B_*(*t*_0_) from Equation 2. Thus the figure illustrates the relative numbers of ultimately transmitted mutations that occur at each time during the infection time course.

## DISCUSSION

We develop a model of within-host pathogen evolution, and use this to predict the fate of *de novo* mutations that occur during disease transmission cycles. Using parameter values specific to influenza A virus and an estimate of the non-lethal mutation rate for IAV, our results predict that the probability that at least one copy of a *de novo* nucleotide substitution is transmitted to the subsequent host is about 5 × 10^−4^ per substitution per site, assuming the mutation is either neutral or beneficial. Multiplying by three possible nucleotide changes and the ≈ 13,600 sites in the IAV genome yields an estimate that as many as 20 sites in the founding dose for the recipient may contain substitutions that occurred *de novo* in the donor. This upper bound, however, must be corrected by the fraction of non-lethal mutations that are either neutral or beneficial. If approximately half of all non-lethal mutations are neutral or beneficial, as reported for another single-stranded RNA animal virus with a similar genome size (Sanjuan *et al*. 2004), we predict each recipient founding dose will contain about ten *de novo* substitutions. If the fraction of neutral or beneficial mutations, among non-lethal mutations, is closer to 10% (see Eyre-Walker and Keightley (2007) for review), we predict about two new substitutions per transmission. As demonstrated in Figures S2 and S7, these estimates scale directly with the underlying mutation rate and the size of the transmission bottleneck. Despite this inherent uncertainty, our results predict that a small handful of mutations occurring *de novo* in the donor will be transmitted to each recipient of IAV.

Our approach makes the simplifying assumption that the founding infectious dose in the donor is phenotypically, but not genetically uniform. Thus the predicted *de novo* mutations may occur on different genetic backgrounds circulating within the donor. Recent evidence suggests that multiple lineages are transmitted between donor-recipient pairs in IAV (Poün *et al*. 2016), and it seems unlikely that all transmitted lineages would be phenotypically identical. Thus a clear direction for future work would be to expand our approach to track multiple distinct lineages within the host, and predict the fates of mutations occurring on these backgrounds.

We can also take our estimate of (1.5 × 10^−3^ non-lethal substitutions per site per transmission event) × (10-50% neutral or beneficial) to predict 1.5 — 7.5 × 10^−4^ substitutions per site per transmission event. These values are consistent with the observed evolutionary rate of IAV throughout a seasonal epidemic, 2× 10^−3^ substitutions per site per year in the nonstructural (NS) gene (Kawaoka *et al*. 1998), if the chain of influenza transmission typically involves 3 to 13 transmission events per season.

Although transmission bottlenecks in IAV, as in many other pathogens, can be extremely severe, our results are consistent with previous work demonstrating that the period of growth between population bottlenecks has an even greater impact (Wahl *et al*. 2002); this period of sustained population expansion promotes the survival of new mutations, as seen more generally in any growing population (Otto and Whitlock 1997). The rapid growth of influenza during early infection, from a relatively small infectious dose to peak viral loads many orders of magnitude larger, implies that neutral substitutions, or mutations conferring even a small benefit, will have ample opportunity to compete with founder strains. This further implies that the life history of influenza A should be well adapted to the disease transmission cycle in humans, in other words, selection has the opportunity to rapidly fine-tune the life histories of pathogens experiencing extreme transmission bottlenecks.

This result is consistent with previous theoretical (Bergstrom *et al*. 1999) and experimental work on viral evolution (Duarte *et al*. 1992; Duarte *et al*. 1993; Novella *et al*. 1995; Novella *et al*. 1996). The latter work focused on the loss of fitness due to population bottlenecks, but fitness could be maintained or improved when the bottleneck size was as large as five or ten individuals (Novella *et al*. 1996). Similarly, Bergstrom *et al*. 1999 predicted that viral pathogens would be well-adapted if the bottleneck size is large (or order five or ten), and the number of generations between bottlenecks is large (of order 25 or 50). The parameter values we explored for influenza A correspond to over 25 population doublings between transmission events, with bottleneck sizes of 10 to 200, and are thus consistent with a parameter regime in which the pathogen is able to improve or maintain fitness.

The use of a specific life-history model imposes natural limits on the growth rate and thus the selective advantage that can be achieved by budding viruses. For the parameters specific to influenza A, changes to the clearance rate of the free virus or death rate of infected cells could only achieve a selective advantage of *s* < 0.1. This occurs mathematically because these rates cannot be reduced below zero; it follows intuitively because even if infected cells or virus never die or lose infectivity, growth remains limited by other processes. Mutations with larger beneficial effects, in the range 0.1 < *s* < 0.2, are accessible only by reducing the eclipse phase, or through very large magnitude changes to the attachment or budding rates. Given that predicted differences in survival probability for the different traits are rather modest (Figure 5), these results suggest that small magnitude changes in the eclipse timing of influenza A will be subject to selective pressure. The limits we observe in the achievable growth rate suggest that larger effect beneficial mutations in influenza A are not only unlikely, they may not be physically possible given the life history of this virus.

We have focused this study on the in-host life history of the virus. In principle, however, a beneficial mutation could also affect the transmissibility of the lineage (parameter F), producing virions that are preferentially transferred to a new host (Handel and Bennett 2008). This would be distinct from mutations that increase viral load; mutations affecting F would increase the probability that an individual viral particle is transmitted, for example by prolonging the stability of the virion in the external environment.

These results explore mutations affecting a single trait in isolation. Clearly higher fitness could be achieved by mutations that affect several traits, if beneficial pleiotropic mutations are available. Previous work suggests that the survival probability of pleiotropic mutations typically falls between the predictions obtained for single-trait mutations of equivalent selective effect (Wahl and Zhu 2015). In addition, we have investigated the transmission of *de novo* mutations when rare. Given the magnitude of the viral loads measured in influenza A, it is clear that multiple beneficial mutations could emerge and compete before the virus is transmitted to a new host. Thus we would expect that clonal interference and multiple mutation dynamics might come into play in describing the adaptive trajectory more fully (Desai and Fisher 2007; Desai *et al*. 2007).

A limitation of the model is that the immune response is not explicitly included as a dynamic variable. Innate immunity is activated when an infection is detected, which is usually within the first few hours of infection. Adaptive immunity, however, is activated, at the earliest, three days post-infection (Tamura and Kurata 2004). Since our model addresses early infection (up to 54.5 hours post-infection) adaptive immune effects are assumed negligible. The innate immune response, however, cannot be neglected, as its main purpose is to limit viral replication (van de Sandt *et al*. 2012). In our approach, innate immune mechanisms are included in the viral clearance and infected cell death rates, but are assumed to be constant throughout this early stage of the infection. This phenomenon has been reviewed in some detail in previous work (Smith and Perelson 2011; Boianelli *et al*. 2015; Baccam *et al*. 2006; Beauchemin and Handel 2011), from which it is clear that directly incorporating the immune response is necessary for an accurate representation of the full time course of infection (Boianelli *et al*. 2015). Even when limiting our attention to early infection only, interferon-I and natural killer cells could be included to more accurately model innate immunity (Boianelli *et al*. 2015). However, the complexity of the immune system creates a significant challenge in accurately modeling influenza A dynamics, even during this initial time period (Boianelli *et al*. 2015). In particular, many key parameters of immune kinetics remain unquantified, creating additional uncertainty (Dobrovolny *et al*. 2013).

Finally, it is well understood that antigenic drift is associated with the evolution of influenza A virus (Carrat and Flahault 2007). Antigenic drift would be formalized in our model as a reduction in the death rate of infected cells or the clearance rate of free virions, as these life history parameters would be improved by any immune evasion. In fact, Figure 2 predicts that for mutations with small selective effects (*s* < 0.08), of all possible mutations with the same selective effect, clearance mutations are the most likely to survive when rare. Thus mutations affecting the viral clearance rate are most likely to adapt. This could shed light on the mechanisms underlying the maintenance of antigenic drift, however much remains to be understood about the complex transmission and evolutionary dynamics of influenza A virus. It is our hope that predicting the fate of *de novo* mutations affecting IAV life history is an important piece of this interesting puzzle.

## ACKNOWLEDGMENTS

J.N.S.R. was funded by the Ontario Graduate Scholarship (OGS). L.M.W. is funded by the Natural Sciences and Engineering Research Council of Canada (NSERC). We thank Catherine Beauchemin for a number of insightful and helpful comments.

## APPENDIX

Let *p_lmn_*(*t*) be the probability that *l* free virions, *m* infected cells, and *n* mature cells exist in the focal lineage at time *t*, and let *A*(*t*) denote the time-dependent per virion attachment rate, which depends on the available target cells, *y_T_*(*t*), as predicted in the deterministic model 1. Parameters *B, C*, and *D* represent the budding, clearance and cell death rates, while *E* denotes the rate at which cells exit the eclipse phase and begin budding. Although the stochastic model follows the mutant lineage, for simplicity we will use *A* as opposed to *Ã*, etc., throughout the Appendix. Also for notational clarity we illustrate the case *k* = 1. Taking into account the stochastic events of attachment, budding, clearance, cell death and cell maturation, is is straightforward to demonstrate that the probability generation function (pgf) describing the time evolution of the lineage must satisfy:

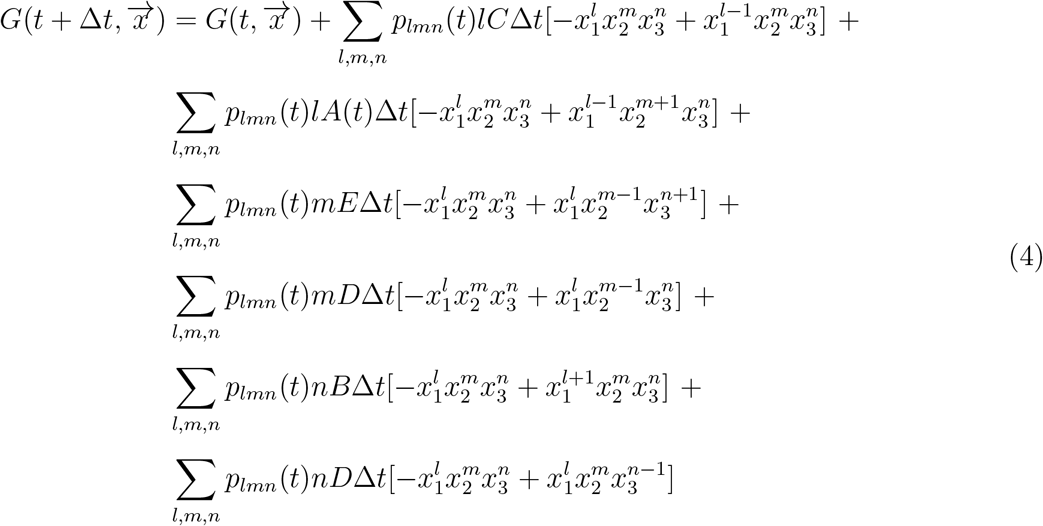

Taking the limit as Δ*t* → 0, Equation 4 yields the following linear partial differential equation:

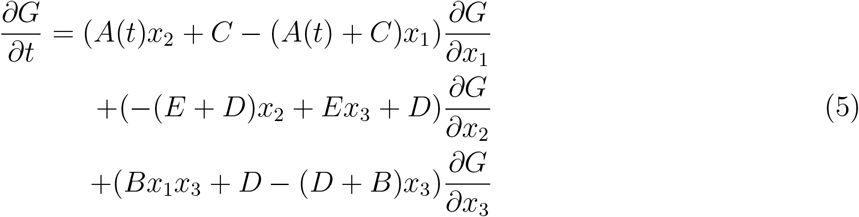

Equation 5 can be converted to a system of ordinary differential equations using the standard method of characteristics, which yields the following system of ordinary differential equations

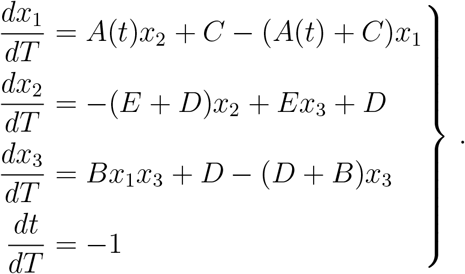

This system can be solved numerically to determine the value of *G* at time *τ*, given the known initial condition corresponding to a single free virion at time *t*_0_, *G*(*t*_0_, *x*_1_, *x*_2_, *x*_3_) = *x*_1_. For convenience we let 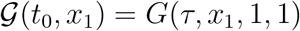 under this initial condition.

The function 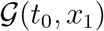, then, gives the distribution of free virions at time *τ*, just before disease transmission, given the lineage began with a single virion at time *t*_0_. Composing this with the pgf of the bottleneck process, we obtain 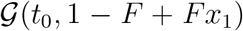 as the pgf describing the distribution of free virions transmitted to a new host (given that one new host is infected). The probability that a given lineage, that arose at time *t*_0_, is *n*ot transmitted to the new host is obtained by evaluating at *x*_1_ = 0:

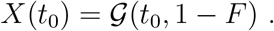

We then use this to compute the expected rate at which surviving mutant strains appear at time *t*_0_, where “surviving” means the lineage will be transfered to the next host:

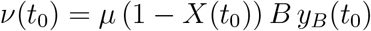

where *μ* is the probability that the mutation of interest occurs, per new virion produced. We use this to compute 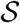, the expected number of times that the mutation of interest occurs *de novo*, over the course of the infection, and survives to be transmitted to the next host:

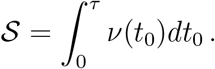

Consider dividing the time interval (0, *τ*) such that *δt* = *τ*/*N* and *t_i_* = *iδt*. In this case for small *δt*, the quantity *ν*(*t*_0_)*δt* approximates the probability that a surviving mutation occurs during time interval (*t*_0_, *t*_0_ + *δt*). This allows us to compute 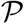, the probability that the mutation of interest occurs *de novo* during the course of the infection and is transmitted to the new host:

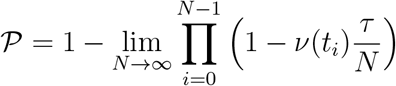

which by product integration can be succinctly expressed as:

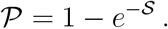

We also compute the expected number of mutant virions transmitted to the recipient host, 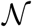. We do this by first computing 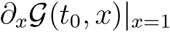, which gives the expected number of mutant virions at time *τ*, given that a mutant virion was produced at time *t*_0_. We multiply this value by the number of mutant virions being produced at time *t*_0_, *μBy_B_*(*t*_0_), and integrate from 0 to *τ*, to get the total expected number of mutant virions at time *τ*. Multiplying by the bottleneck fraction, *F*, gives the expected number of mutant virions transmitted to the recipient host:

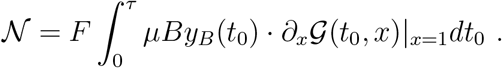

Note that 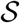 and 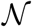 differ because each *de novo* mutation produces a lineage that could in principle contribute more than one virion to the recipient.

## Supplementary Figures

**S1. Expected number of successful** *de novo* **occurrences,** 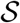

**Figure S1:**
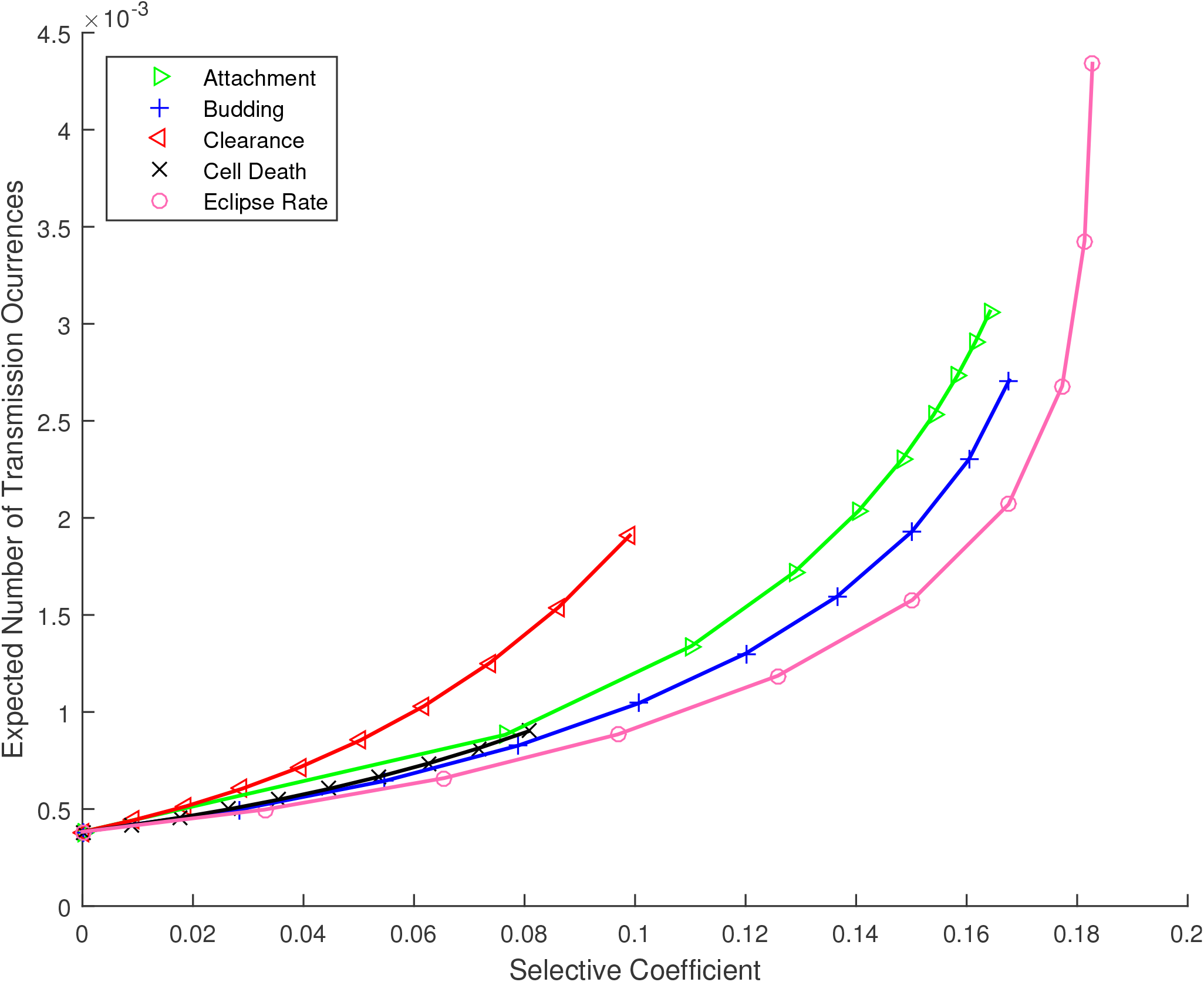
The number of times that a given mutation is expected to arise *de novo*, during a single infection time course, and produce a lineage that is transmitted to the next infected individual (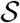, as described in the Appendix), versus the selective coefficient. This quantity scales linearly with μ, the mutation rate.

**S2. Expected number of transmitted mutant virions,** 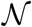

**Figure S2:**
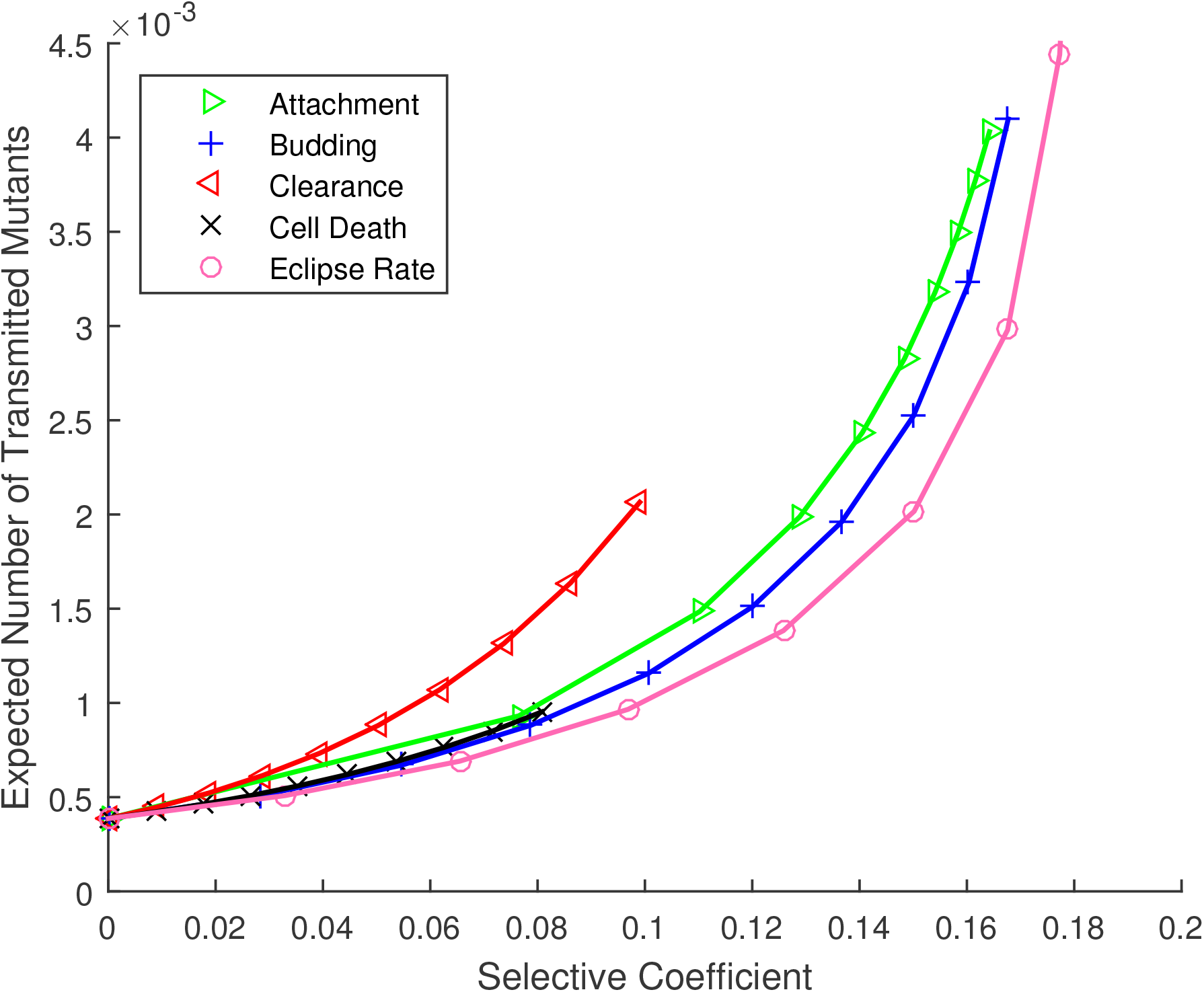
The expected number of virions transmitted to the recipient that have arisen *de novo* during a single infection time course in the donor (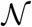, as described in the Appendix), versus the selective coefficient.

**S3. Effect of non-exponential cell lifetimes: Deterministic time course**

**Figure S3:**
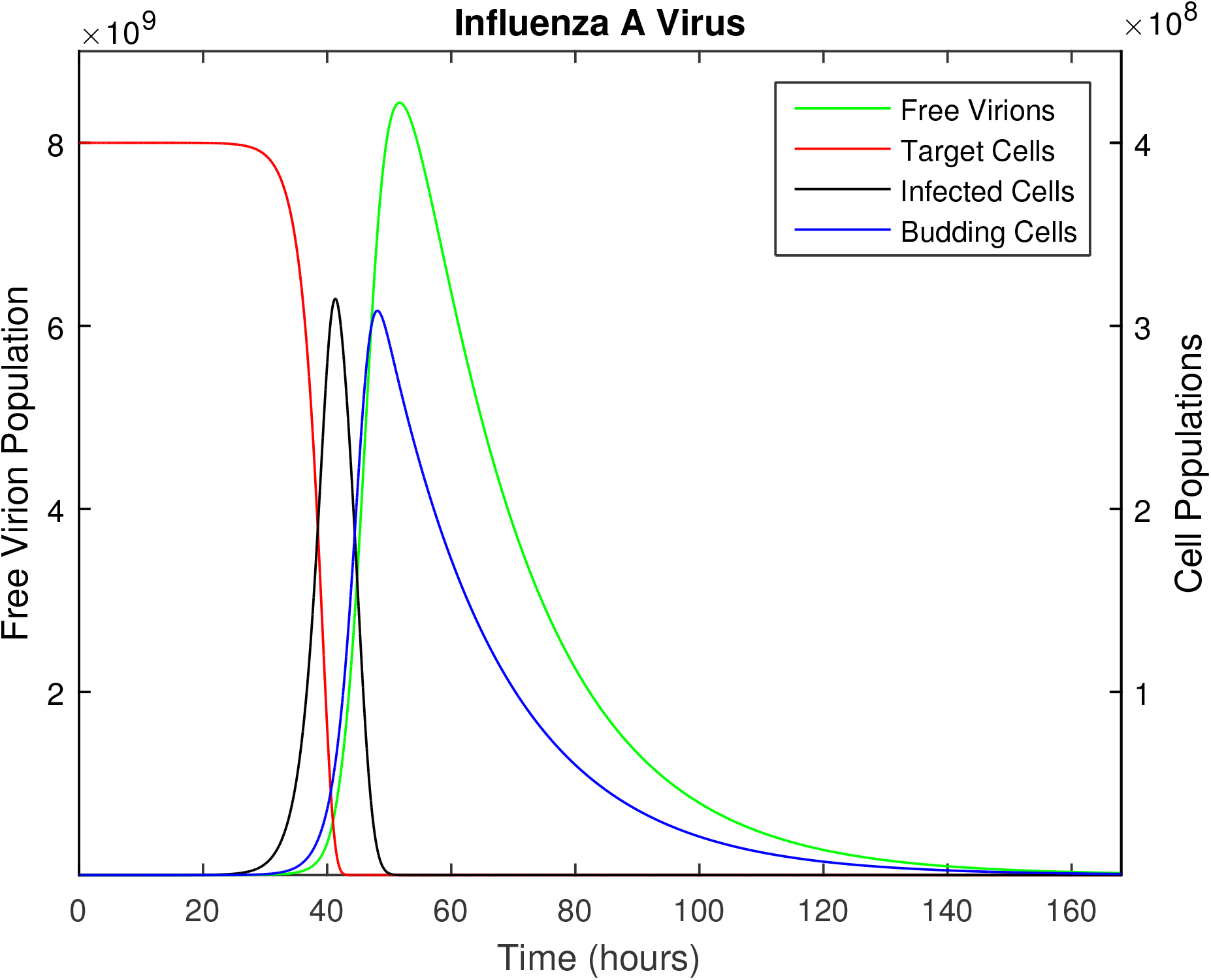
The time course of influenza A infection over the span of a week (168 hours). This figure is analogous to Figure 1, except that the cell death rate, *D*, has been set to zero during the eclipse stages, and increased during the budding phase such that the mean infected cell lifetime is unchanged. Other parameters as provided in Table 1. The infection time course is relatively insensitive to these changes in the distribution of infected cell lifetimes.

**S4. Effect of non-exponential cell lifetimes: Transmission probability**

**Figure S4:**
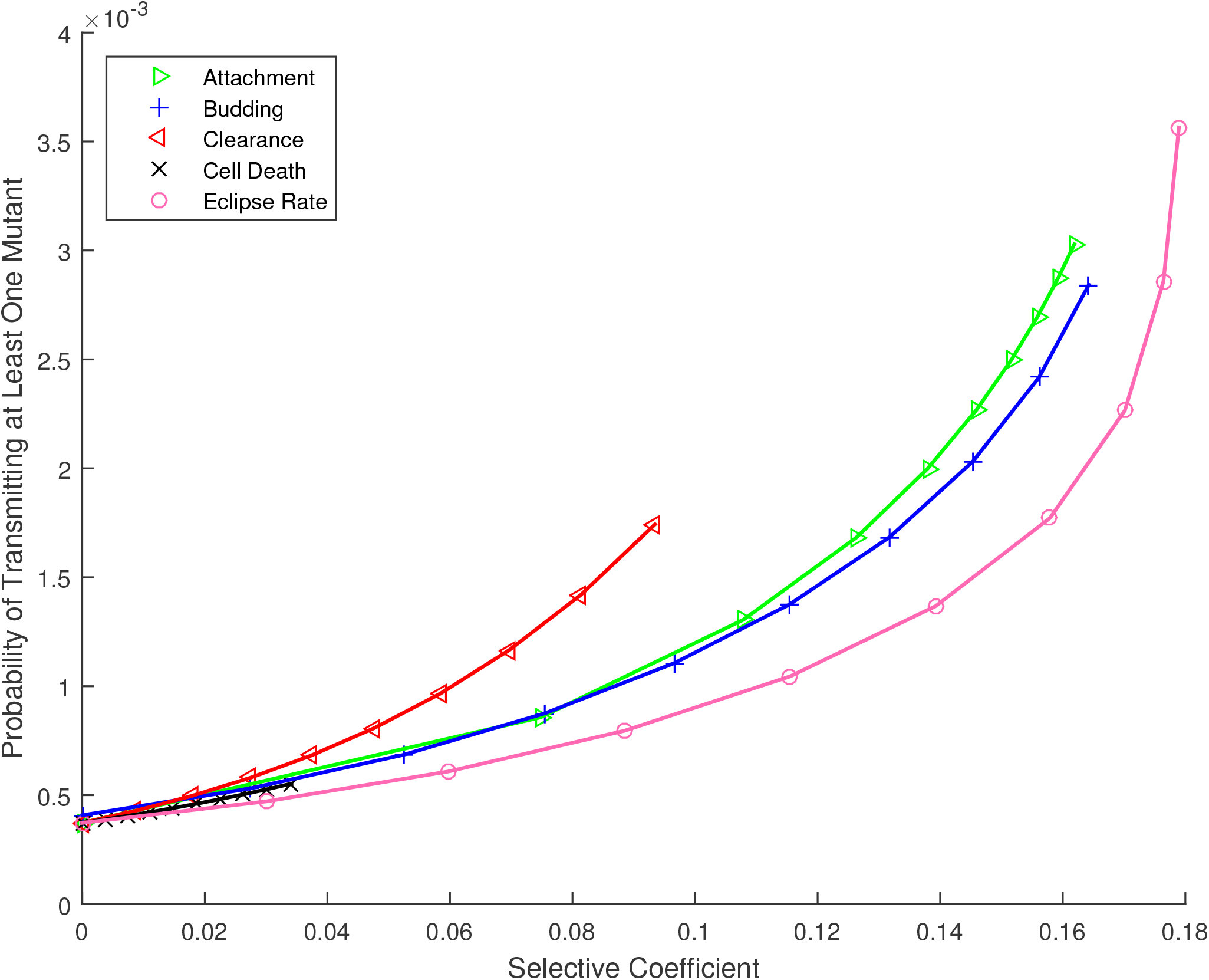
The probability that at least one *de novo* mutation arises during the infection time course and is passed to the next host, 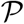, versus the selective coefficient, *s*. This figure is analogous to Figure 2, except that the cell death rate, *D*, has been set to zero during the eclipse stages, and increased during the budding phase to yield the same mean infected cell lifetime. Other parameters as provided in Table 1. We find that the transmission of *de novo* mutations is insensitive to these changes in the distribution of infected cell lifetimes.

**S5. Effect of available target cells: Deterministic time course**

**Figure S5:**
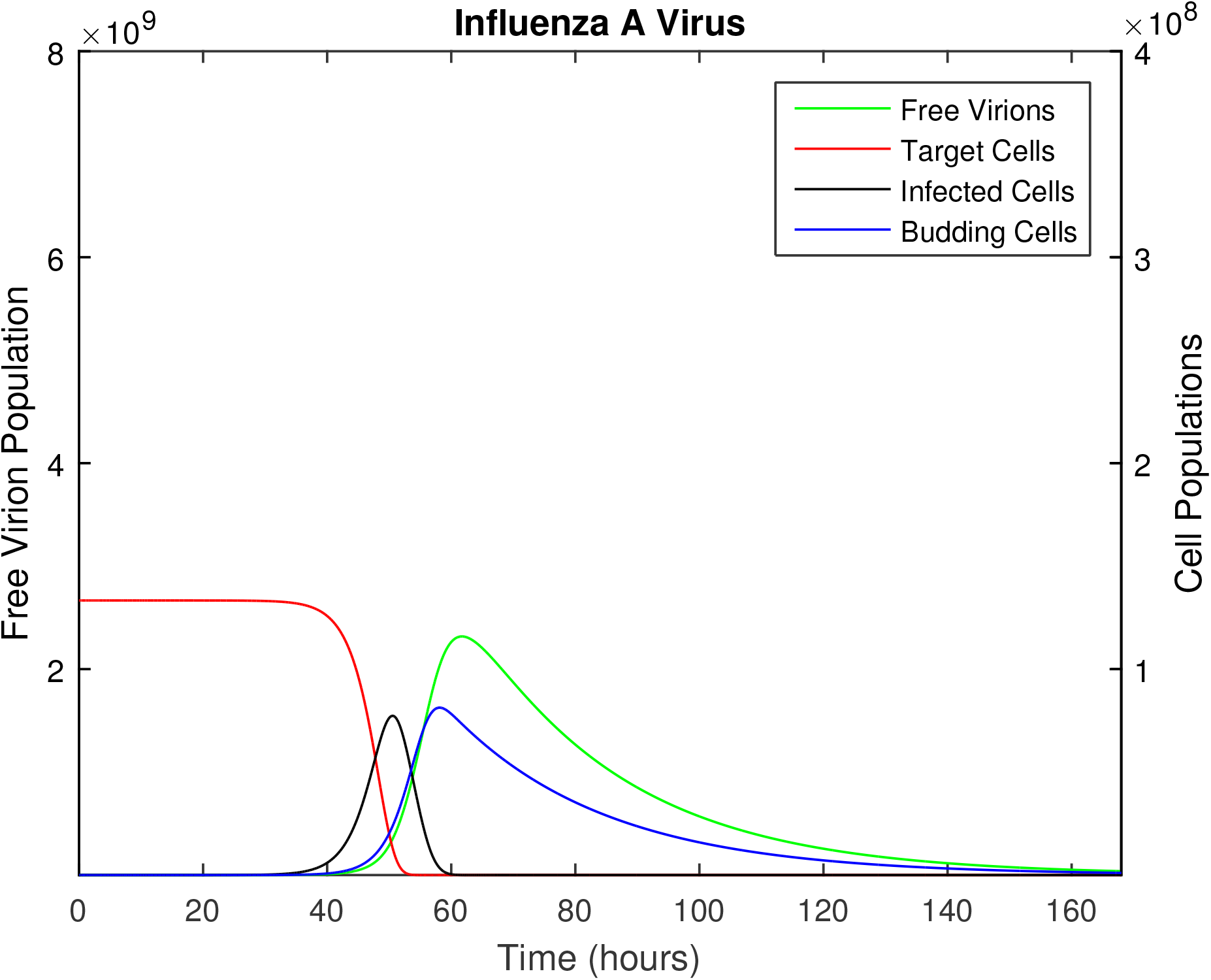
The time course of influenza A infection over the span of a week (168 hours). This figure is analogous to Figure 1, except that the initial target cell population, *y_T_*(0), has been reduced by a factor of 3. Although complete desquamation is the expected outcome of the infection, it is possible that spatial considerations might spare a fraction of the epithelial cells in the upper respiratory tract; we therefore included this case in sensitivity analysis. Other parameters as provided in Table 1. Comparing with Figure 1, the magnitude of the infection is scaled and the dynamics are slightly delayed. However this has little impact on the probability of transmission of a mutation (see Figure S6).

**S6. Effect of number of available target cells: Transmission probability**

**Figure S6:**
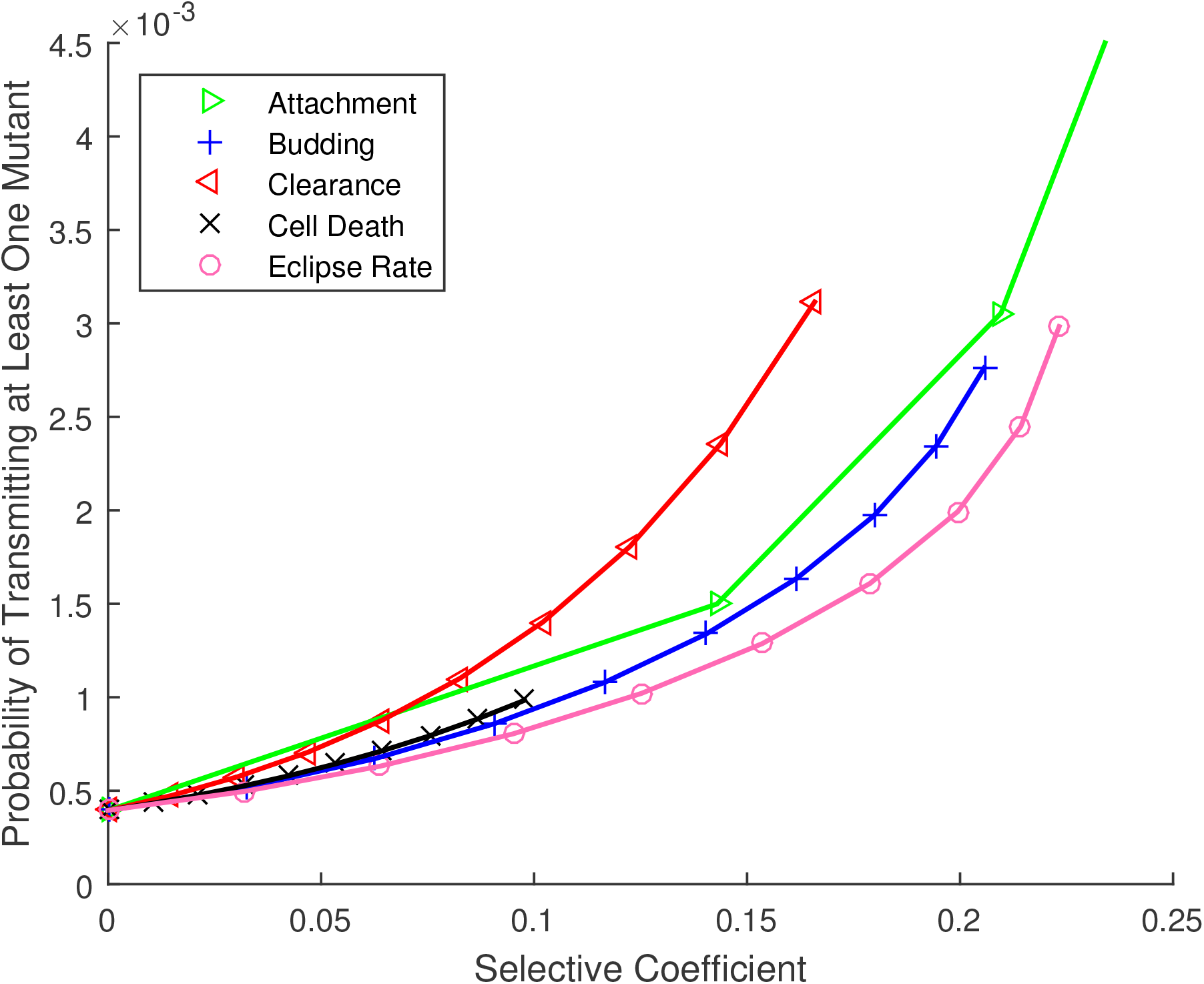
The probability that at least one *de novo* mutation arises during the infection time course and is passed to the next host, 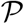, versus the selective coefficient, *s*. This figure is analogous to Figure 2, except that the initial target cell population, *y_T_*(0), has been reduced by a factor of 3. Other parameters are as provided in Table 1. We find that the transmission of *de novo* mutations is insensitive to the initial number of available target cells.

**S7. Effect of varying mutation rate**

**Figure S7:**
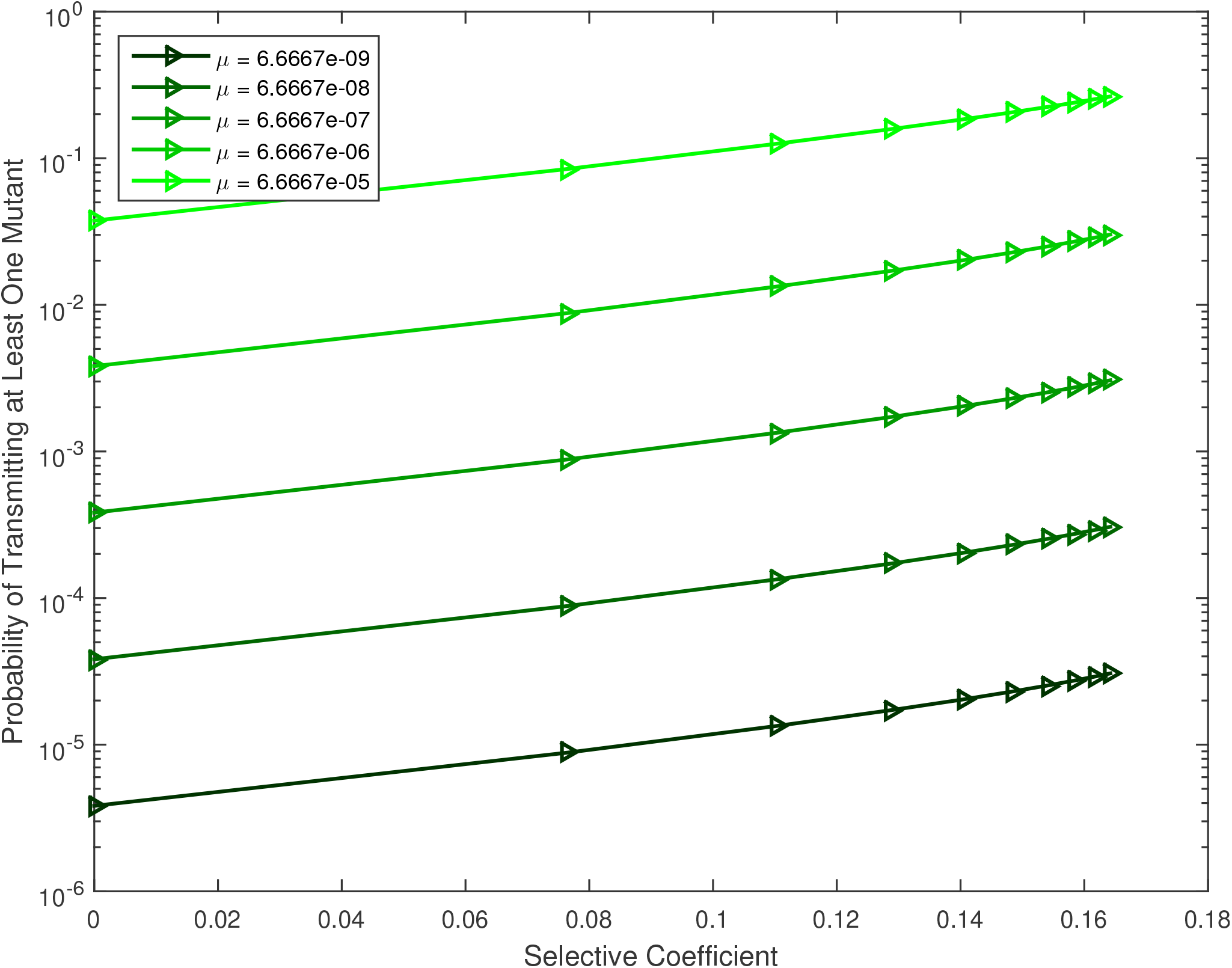
The effect of mutation rate on the probability of transmission. The probability that at least one copy of a *de novo* mutation is transmitted to the next host is plotted against the selective coefficient, for a mutation that increases the viral attachment rate. The mutation rate per replication event, *μ*, is varied. Other parameters as provided in Table 1.

